# Using feedback in pooled experiments augmented with imputation for high genotyping accuracy at reduced cost

**DOI:** 10.1101/2023.12.12.571203

**Authors:** Camille Clouard, Carl Nettelblad

**Affiliations:** Division of Scientific Computing, Department of Information Technology, Uppsala University, Uppsala, Sweden; SciLifeLab, Science for Life Laboratory, Uppsala, Sweden

**Keywords:** SNP array, pooling, imputation, iterative refinement

## Abstract

Conducting genomic selection in plant breeding programs can substantially speed up the development of new varieties. Genomic selection provides more reliable insights when it is based on dense marker data, in which the rare variants can be particularly informative while they are delicate to capture with sufficient statistical power. Despite the release of new performing technologies, the cost of large-scale genotyping remains a major limitation to the implementation of genomic selection. We suggest to combine pooled genotyping with population-based imputation as a cost-effective computational strategy for genotyping SNPs. Pooling saves genotyping tests and has proven to accurately capture the rare variants that are usually missed by imputation. In this study, we investigate an extension to our joint model of pooling and imputation via iterative coupling. In each iteration, the imputed genotype probabilities serve as feedback input for rectifying the decoded data, before running a new imputation in these adjusted data. Such flexible set up indirectly imposes consistency between the imputed genotypes and the pooled observations. We demonstrate that repeated cycles of feedback can take full advantage of the strengths in both pooling and imputation. The iterations improve greatly upon the initial genotype predictions, achieving very high genotype accuracy for both low and high frequency variants. We enhance the average concordance from 94.5% to 98.4% at a very limited computational cost and without requiring any additional genotype testing. We believe that these results could be of interest for plant breeders and crop scientists.

**Author summary:** In applications such as large-scale population surveys or plant breeding, the cost of genetic testing can limit the number of samples that are genotyped, or force the reduction to more cost-effective low-density marker panels. A reduction in the number of samples or the number of variants surveyed can reduce the power to detect important genetic correlations. We propose a scheme of pooled genotype testing, which would allow for using half the number of test assays for the same number of individuals surveyed. The data from overlapping pool tests is augmented with genotype imputation. We have previously shown that this approach was competitive, but with some drawbacks. Most strikingly, the error rate for common variants could be in the range of 10%. Now, we propose a new computational method for reconstructing SNP genotypes with pooling and imputation, adding an iterative coupled model connecting the two. This model allows us to exploit the advantages of both methods and achieves consistently high genotype reconstruction accuracy. We demonstrate the performance of our approach on a hypothetical plant breeding application based on a public genetic dataset from wheat samples. However, main aspects of the methodology would translate to many other settings.

## Introduction

Modern plant breeding programs have increasingly involved genetic data, typically the genotypes of single nucleotide polymorphisms (SNPs), for supporting the selection process and accelerating the development of new varieties. Related methods such as genomic selection (GS) and genome wide association studies (GWAS) are more accurate if using high-density marker data, as well as the quality of the data collected from e.g. SNP-microarrays directly impact the accuracy of the genomic predictions in GS [1–3]. While they are restricted to already known markers and not suitable for variant discovery studies, SNP-microarrays have remained popular for routine genotyping and they provide more accurate measurements [4] than the next generation sequencing (NGS) techniques. GS and GWAS gain in statistical power with larger population sizes, but genotyping more samples at high density also leads to more expensive data collection. Despite their cost-effectiveness, imputation methods for augmenting low-density genotype data to high density have a known weakness at capturing rare variants, that is with a minor allele frequency (MAF) less than 5%, while these variants can have a great significance in GS.

We have previously explored the combination of pooling and imputation by simulating SNP chip pooling experiments based on available genotype data of recombinant inbred lines of bread wheat (*Triticum aestivum*). We found that decoding pooled experiments with a nonadaptive and overlapping design, followed by data augmentation via imputation with a suitable method, can improve the genotyping results for rare variants compared to conventional imputation workflows on less dense arrays. Moreover, our strategy can cut the number of required SNP array plates by half. Earlier studies [4, 5] with human data have proposed similar hybrid approaches for genotyping and investigated the sequencing of overlapping DNA pools in conjunction with imputing genotypes assayed on microarrays, and their findings are similar to ours. Firstly, sequencing pools instead of individual samples reduces the experimental cost for genotyping, for instance the cost for library preparation can be divided by 3 [5]. Secondly, combining the genotypes decoded from the pooled sequenced reads to the imputed data results in highly accurate genotyping for SNPs with both low and high allele frequency. The pooling scheme takes advantage of the sparsity of the variants with low frequency that are usually missed by the imputation methods. Pooling generates very noisy outcomes for common variants due the information theoretic bounds [4], but these variants can be accurately imputed thanks to models that take into account the correlation structure between different genetic variants at the population level. The key to these combined genotyping approaches was the use of a likelihood framework that has a greater flexibility than a purely combinatorial approach [4, 5]. Such frameworks allows for carrying the information from imputation e.g. the linkage disequilibrium (LD) between consecutive markers as well as the MAF, over to the procedure for decoding the pools. [4] proposed to compute the genotype likelihoods from three different perspectives and take their product as a composite likelihood for calculating the final estimate of each most likely genotype at each entry. [5] studied an iterative refinement of the imputed genotypes with approximate gradient descent based on a maximum a posteriori (MAP) strategy which handles the minor and the major alleles separately. Both studies demonstrate the performance of injecting information from imputation into the decoding procedure and these results have motivated the approach we present here.

This paper explores what gains in genotyping accuracy can be achieved if we enforce consistency between the pooled observations and imputed genotypes in a scheme similar to the one we first proposed in [6]. We suggest an iterative coupling procedure between local pattern-consistent decoding and population-based imputation in a likelihood framework. At every iteration, we use the imputed data as feedback for applying a correction to the decoded genotype likelihoods before passing the corrected likelihoods to another round of imputation. We realize a sizeable gain in accuracy of the reconstructed genotypes through the iterations of our coupled model. We also explore the computational performance of our simulations and experiments. We do not conduct either any detailed quantification of the costs associated to this structure, but we believe that our strategy can in some scenarios be a valuable addition to existing schemes for cost-effective genotype reconstruction at scale, such as non-pooled low-coverage sequencing followed by imputation.

## Materials and methods

We simulate pooled experiments with genotype data of SNPs in an identical fashion to our previously published protocol [6]. We consider the results from our pooling simulation to be experimental observations analyzed in later stages of the workflow. The initial decoding algorithm we use computes estimates of the genotype probabilities at any variant and in each sample via expectation-maximization. That is, this step is agnostic for LD and allele frequencies at the population level since each variant is treated independently of the other ones, and likewise for each pooling block. Coalescence-based imputation, as we conducted in a second step, is expected to indirectly enrich the genotype probabilities estimated from the pools with information about the global genetic structure in the population. Imputation reconstructs the sequence of genotypes for each sample as a mosaic of the available template haplotypes, which enforces similarity of the genetic profiles across the study population and the reference panel. Though, there is no guarantee that the imputed genotypes are consistent with the decoded prior genotypes passed as input. Our idea is to iteratively apply point-wise corrections to the decoded genotype probabilities based on the imputed results, in order to softly achieve consistency between the decoded outcomes at the local level of the pools, and the predictions from imputation at the global population level. Such sequential feedback schemes between two models have been implemented in various other research fields. A diagram of our iterative procedure is presented in Fig. 1.

**Fig 1.**
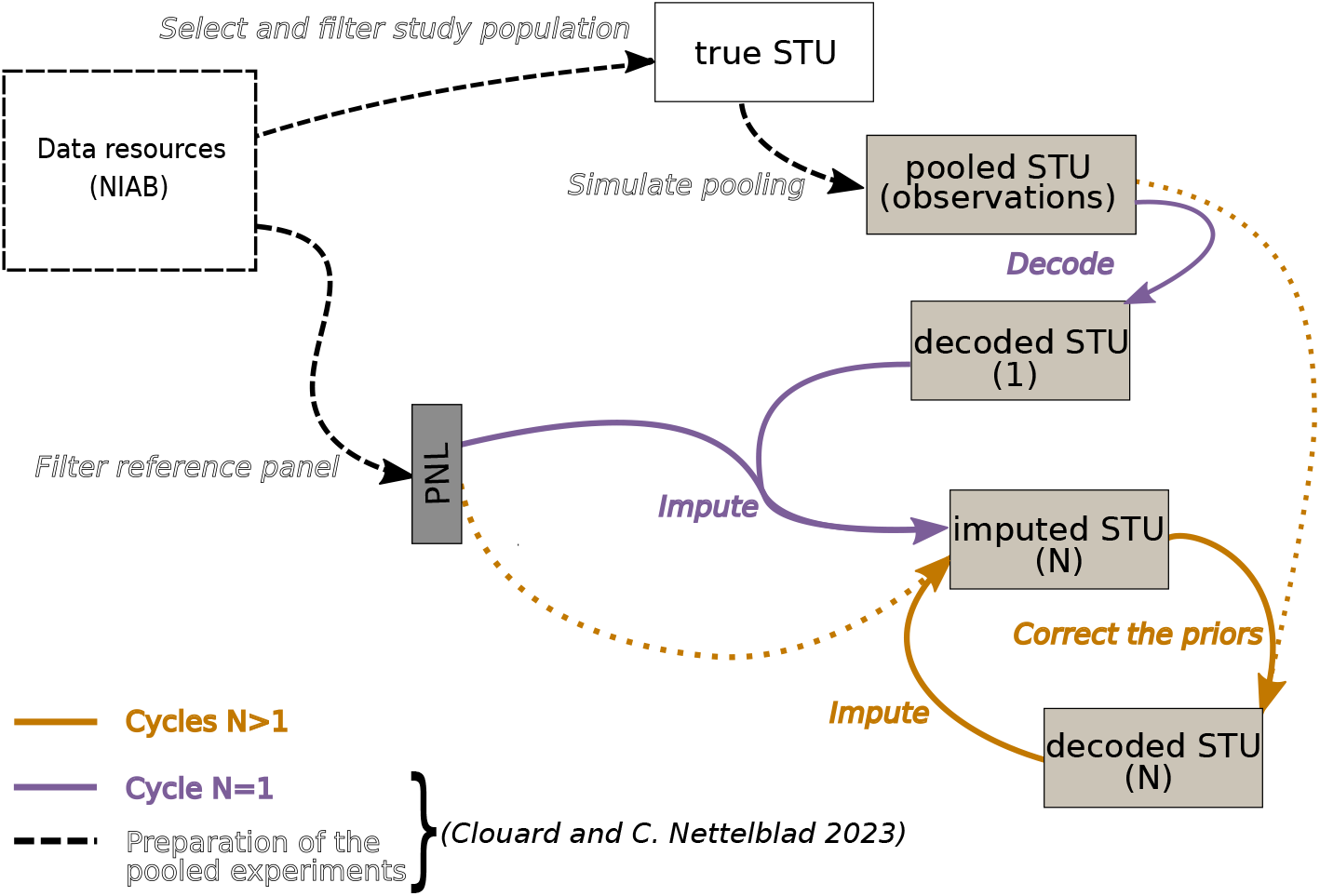
Experimental steps of the iterative genotype pooling and imputation strategy. The data resources used are provided in open access by UCL and were developed by the National Institute for Applied Botany (NIAB) [7]. They consist in a mapping population named *NIAB Diverse MAGIC wheat population* for one thing, and of the set of 16 founders from which the inbred lines were derived. We use the set of founders as reference panel and randomly select 496 samples in the mapping population to serve as study population for our pooling and imputation simulations. The preparation of the pooled experiments and the cycle *N* = 1 are executed with a workflow we developed [6]. The reference panel is assumed to be individually fully genotyped while the study population is mixed into pools previously to full genotyping of the pools. The pooled genotyping results are decoded with a statistical inference method to individual genotype likelihoods, which are in their turn passed to our imputation algorithm. The later cycles *N >* 1 are the focus of the research in this manuscript. We want to investigate whether the additional iterations of correction of the decoded genotype likelihoods, followed by imputation from the corrected values, can improve the overall accuracy of genotyping in the study population (*imputed STU (N)*) with our pooling strategy. The correction of the priors in the decoded data from the cycles *N* to *N* + 1 is based on a comparison between the likelihood of detecting each allele in the pools after the initial pooling (*N* = 1), and likewise in the simulated the pools from the imputed data in cycle *N* . Such a mechanism enables for incorporating extrinsic information forwarded through the population-based imputation model into the decoding model which is otherwise local to each variant and each pooling block.

### Experimental data

The cohort of 496 samples used as study population in imputation is part of the *NIAB Diverse MAGIC wheat* population [7] of inbred lines, and the 16 founders of this experimental population form the reference panel. The genotype data we are looking at, both in the study population and in the reference panel, were originally obtained. with sequencing technologies. The authors of the dataset applied strong filtering of the variants that were sequenced, active removal of heterozygous data, and imputation of the missing genotypes in the inbred lines [7], such that we have considered that the remaining SNPs form a set of markers that could be used, after proper testing, for developing a microarray. Therefore, we have treated the genotype data as if would derive from a hypothetical SNP-chip of 55K markers, and we have focused on the loci on the chromosome 1. While bread wheat is formally allohexaploid, it can typically be modelled as a diploid-like species [8] due to its subgenome structure that consists in hybridized diploid subgenomes, ignoring rare recombinations between them. After additional filtering of the set of markers present on chromosome 1A, the study and reference data sets comprise 1170 bi-allelic positions.

### Initial round of pooling and imputation

The initial cycle (*N* = 1) consists of a first step of pattern-consistent decoding of the observations at any marker into individual genotype likelihoods, which are denoted with the vector *y*^(1)^. In this first decoding step, we assume error-free genotyping, that is the alleles in the pools are detected with full certainty. In a second step, the genotype likelihoods in *y*^(1)^ are normalized and used as prior genotype probabilities by a population-based imputation method called *prophaser* [9], which predicts for each variant the vector of genotype probabilities *z*^(1)^. For details about the algorithm implemented for pattern-consistency decoding or about the imputation method, and for a detailed analysis of the imputation performance on pooled data, we refer to earlier work [6]. We found that the common variants were imputed with far lower accuracy than rare ones, when measured in terms of concordance and cross-entropy, and we also observed a larger inter-marker variance for these metrics at higher MAF.

### Subsequent cycles of decoding and imputation

The strategy for carrying out the next cycles *N >* 1 is to verify whether the imputed data in cycle *N* are consistent with the initial measurements for the simulated pooled observations. This realizes a coupling mechanism between the model used for pool-local decoding, and the population-based imputation model. If an inconsistency is found, the decoded genotype likelihoods of each sample in the pool used in cycle *N* are corrected, and constitute the values used as prior genotype probabilities to imputation in cycle *N* + 1. Consistency is not evaluated on a per-individual basis nor on genotype matches, but from the alleles that are expected to be detected in the pools simulated from the imputed data. That is, imputed data in a pool is consistent with the observations if the same alleles are detected in both cases. Whenever an allele is not detected in the imputed data whereas it should be observed, a correction is applied to the decoded genotype probabilities. Thus, the decoded genotype probabilities of the samples showing consistent imputed data are not updated at the level of the pool considered. However, because of the overlapping design used, the decoded data in a consistent pool might be changed between consecutive cycles if some samples are involved in other pools that are inconsistent. An example of such case is presented in Supplemental file 1.

### “Repooling”

We use a modified version of our original decoding algorithm used in cycle *N* = 1, that we call *repool*. The modified algorithm compares in each cycle the likelihoods of detecting the alleles for a given variant in the imputed data against what alleles should be detected according to the simulated pools. For instance, an observed pooled genotype value equal to 0 means that the probability of detecting the allele *A*_0_ = 0 in the pool is 1.0, and the probability of detecting the allele *A*_1_ = 1 is 0.0. If the initial genotype of a pool is assayed as equal to 1, both the probability of detecting the allele *A*_0_ and the probability of detecting allele *A*_1_ are equal to 1.0.

We denote any sequence of enumerated genotype data *x*, the decoded data in cycle *N y*^(*N*)^, and the imputed data in cycle *N z*^(*N*)^. *x* is represented as a vector of integer genotype values in *{*0, 1, 2*}, y*^(*N*)^ as a vector of log-genotype likelihoods, and *z*^(*N*)^ as a vector of predicted genotype probabilities. The procedure applied for point-wise correction of the decoded outcomes in cycle *N >* 1 in a single pooling block for a single variant is described in Algorithm 1. For a detailed example of calculations with this algorithm, we refer the reader to Supplemental file 1.

In the first part of the algorithm on line 2-12, the probability of detecting the allele *A* ∈ *{A*_0_, *A*_*1*_ } in the pool *p* from the imputed data 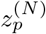 is computed in two steps that proceed from applying the likelihood theory and considering the imputed genotypes for each individual to be independent. First, the probability of any enumerated sequence of genotypes 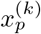 is the product of the individual imputed genotype probabilities. Second, the likelihood of detecting any allele in each enumerated sequence is determined as the allele contribution from the composite genotype of 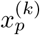, weighted by the likelihood of the sequence. Among the enumerated sequences, some of them might be inconsistent with the observations, as the two examples that are provided in Supplementary file 1. It is important to note here that computing the likelihood of detecting an allele is not the same as computing the expected allelic dosage from imputed data. In our algorithm, both homozygotes for the allele *A* and heterozygotes are assumed to equally contribute to detecting the allele *A* in a pool. The second part of the algorithm on lines 13-22 performs an evaluation of the consistency for each allele separately, and, if necessary, performs a correction for the decoded genotype likelihoods that carry the allele. Likelihoods are only changed to the extent that the imputed likelihoods 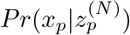 imply inconsistency with the observation in terms of what alleles would be present. If the data are consistent, no change is made between cycle *N* and *N* + 1 (line 15). It should be noted that consistency is distinct from correctly imputed results. An inconsistent imputation is definitely incorrect for at least one individual in the pool, but a consistent imputation might be incorrect for one or more individuals. If the imputed data in the pool *p* are evaluated as inconsistent with the observation e.g. the allele A is not detected in the imputed samples but should be, the deviation is computed as the entropy of detecting the allele A in the imputed data against detecting it in the observation (line 18). In other words, the deviation is proportional to 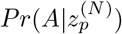. The higher the probability of detection 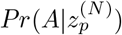 is, the smaller the deviation. The correction applied to the genotype likelihoods compensates for this deviation, weighted by a dampening factor *w* (line 18). This factor lets us adjust the strength of the correction between individual cycles. Too high values can easily result in overcorrection, since the prior for multiple linked markers in the same individual could be updated in a single cycle, causing an overshoot. A higher dampening factor will make the iterative coupling converge in fewer cycles, but with a lower final accuracy in the predicted genotypes.

### Imputation with *prophaser*

Following the correction of the decoded genotype likelihoods, *prophaser* is executed as in the initial cycle, that is with the same reference panel, the same genetic map, and the same hyperparameters e.g. an effective population size equal to 16.

### Technical implementation of the iterations and computational resources

Data downloading and preprocessing, simulation of pooling, as well as all cycles of decoding and imputation, are executed on a cluster node HP ProLiant SL230s Gen8 that has a memory configuration of 128 GiB and consists of 2 eight-core CPUs (Intel Xeon E5-2660).

We extend our workflow [10] by adding to it a Bash script which consists of chained Slurm jobs with job dependencies, such that a new cycle of decoding and imputation starts only if the previous one has successfully completed for all samples.

#### Algorithm 1 Pseudocode for point-wise reinforcement/correction of the genotype prior probabilities from cycle N to cycle N+1 in 1 pooling block for 1 variant

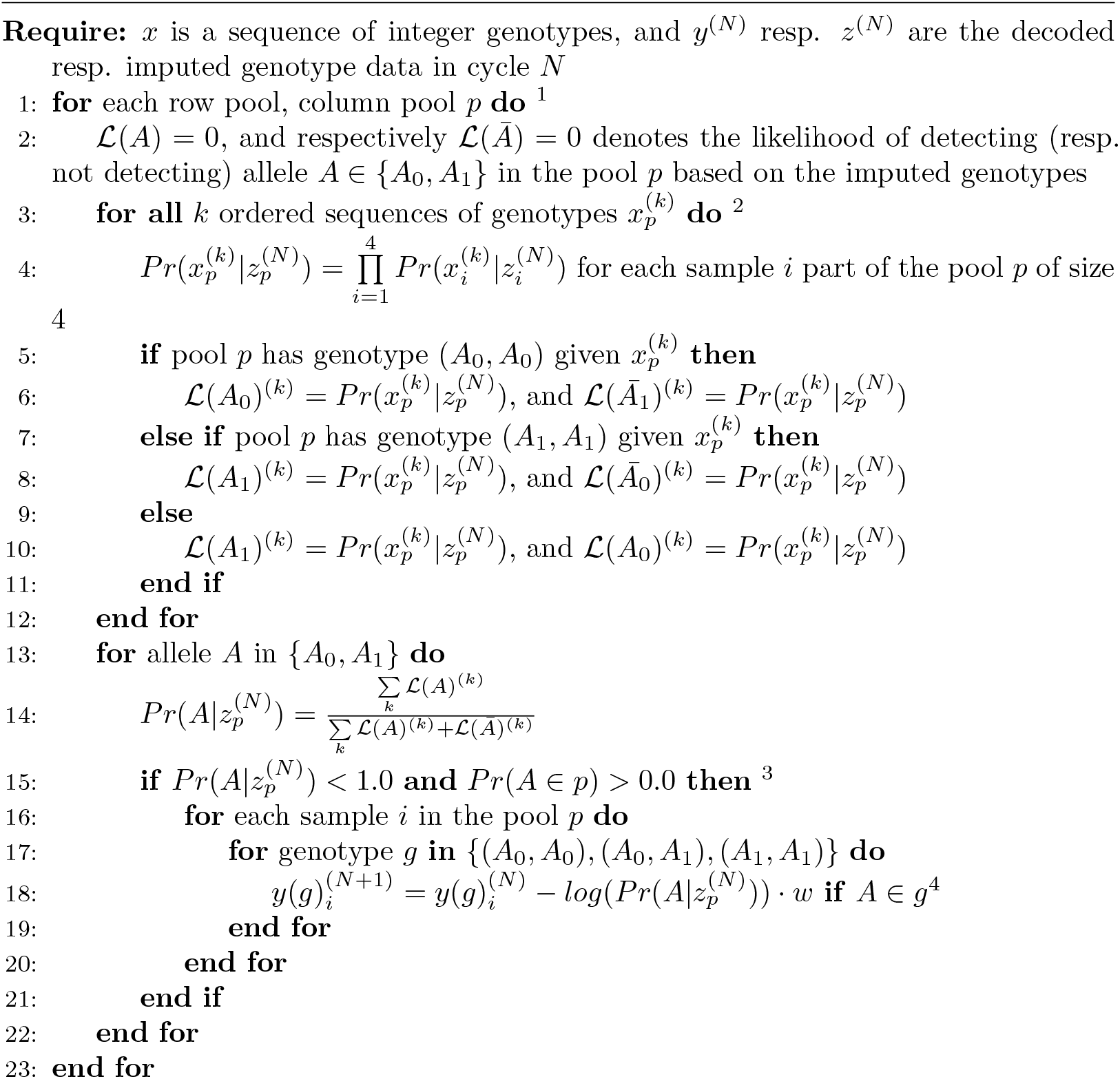

*Notes:*

^1^ Since the pooling design is overlapping and has a degree of intersection equal to 2, each sample is processed twice: once in a row pool, and once in a column pool.

^2^ There are at most 3^4^ sequences to marginalize over in the theoretical case with 1 heterozygous and 2 homozygous genotypes possible for each of the 4 samples in the pool. In the case of the inbred lines of wheat we handle that are fully homozygous, the largest number of sequences is 2^4^.

^3^ If the probability of detecting a given allele in the imputed data is less than 1 but this allele is detected in the observations, the imputation outcome is not considered consistent. The magnitude of the deviation in the imputed data is used in the output step to increase the prior for the missing allele in *y*(*g*)^(*N*+1)^.

^4^ We denote *w* the dampening factor for the pooled outcomes applied at every cycle, and 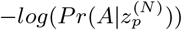 is the correction term. In practice, we apply a max-transformation such that the correction term has a lower bound set to *−log*(10^*−*3^). If 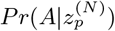 is close to 1.0, the correction applied is almost 0. In other words, the higher the inconsistency between the imputed and the initially decoded data is, the more the decoded genotype likelihoods are corrected.

### Metrics for analyzing the genotyping accuracy and the computational performance

As we are interested in comparing the accuracy of the genotype predictions after any additional cycle against the results obtained in the cycle 1, we use the same metrics as in our first study [6], that is the genotype concordance and the cross-entropy averaged per variant in the study population and presented as the average per binned MAF, with bins of size 0.01. We recall that the genotype concordance measures the degree of similarity between the true and the predicted genotype represented as integers, it is equal to 1 if there is no difference between the true and the predicted value. The cross-entropy renders the element-wise distance between the true and the predicted genotype probabilities, it is equal to 0 if the ground truth and the prediction are identical.

In order to get more detailed insights into what markers are corrected in any cycle, we compute separately the marker-wise genotype concordance and cross-entropy for the the entries with modified priors on the one hand, and for the unchanged data on the other hand. In this case, there is no averaging per variant before averaging per MAF-bin as this would be less relevant given there is a variable number of entries that have a changed prior at each locus. We consider a prior is significantly modified between two cycles if the difference of the prior log-likelihood is larger than *ϵ* = 10^*−*5^.

Last, we measure the time and memory usage for computation, both for the correction of the priors in the full study population, and for the per-sample imputation. Measuring the global execution time for all cycles is less relevant, since the time used for the orchestration of the full set of chained jobs with dependencies will mostly be dependent on the load level of the queuing system.

## Results

The results for the genotyping accuracy we give in this manuscript are computed in the imputed study population after various cycles (*imputed STU (N)* in Fig. 1) w.r.t. the corresponding true genotype data (*true STU* in Fig. 1). In total, 42 cycles are run. The dampening factor *w* for the genotype likelihoods is set to 0.01 in the results presented and discussed, but supplementary results with other values for *w* can be found in Fig. S2.1 to S2.10, and Table S2.1. Based on the criteria of the speed of convergence against the improvements yielded in the genotyping accuracy, we think that a dampening factor *w* = 0.01 offers a good compromise and therefore only present the results for this series of experiments in the main manuscript.

### Impact of additional cycles of correction and imputation on the accuracy of genotyping

In Fig. 2, we show the average concordance and cross-entropy with respect to the MAF after the cycles 1, 2, 3, 12, and 42. We observe an overall improvement in both metrics, which confirms that repeated corrections and imputations increase the genotyping accuracy. For instance, the average concordance rises from 94.5% (N=1) to 98.4% (N=42). We also note that much of the gain in accuracy occurs already in the first few cycles. For instance, the average concordance is improved by almost 1% from cycle 1 to cycle 2 but by barely 0.7% in the 31 latest cycles (from N=12 to N=42). The extent of the gains is highly dependent on the MAF. As shown in Table 1, most loci in the set of variants that we study are common variants as 1094 out of 1170 loci have a *MAF >* 0.1. The accuracy of the genotype predictions for these variants appears to be most improved in Fig. 2 with iterative correction and imputations cycles. For the rare variants, we previously found that the genotyping accuracy is about 99% concordance after one cycle of decoding and imputation [6]. This score is improved with additional cycles of correction and imputation as well, however to a smaller extent than for the common variants. After 42 cycles, the variants with *MAF <* 0.1 are close to be perfectly imputed and the concordance for the variants with the highest *MAF* is improved by nearly 10%.

**Table 1.** Statistics for markers per MAF bin in the population of inbred lines (true data)

|  | 0.00-0.05 | 0.05-0.10 | 0.10-0.20 | 0.20-0.30 | 0.30-0.40 | 0.40-0.50 | Total |
| --- | --- | --- | --- | --- | --- | --- | --- |
| Counts | 76 | 260 | 363 | 221 | 139 | 111 | 1170 |
| Proportions<sup>1</sup> | 0.065 | 0.222 | 0.310 | 0.189 | 0.119 | 0.095 | 1.000 |

<sup>1</sup> SNP proportions per MAF bin with respect to the total number of SNPs in the genetic map.

**Fig 2.**
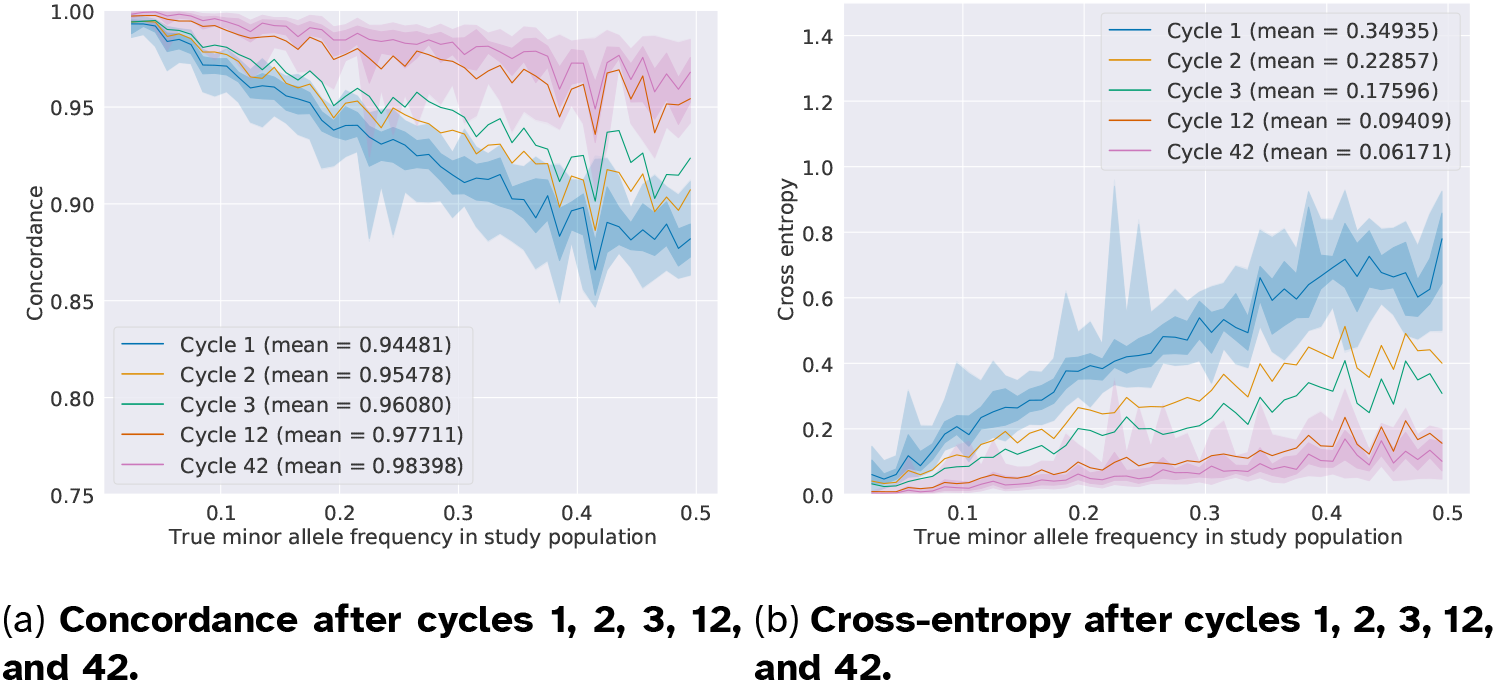
Genotyping accuracy after cycles 1, 2, 3, 12, and 42 (dampening factor *w* = 0.01). The concordance and cross-entropy scores are computed between the imputed data (*imputed STU (N)* in Figure 1) and the filtered data (*true STU*). The imputed markers are sorted per ascending true MAF in the study population and categorized into MAF bins (bin size equal to 0.01). Each marker has a smoothed concordance and cross-entropy score which is calculated as the average value in a rolling window of 5 MAF-consecutive markers. The plain line shows the median accuracy value (concordance or cross-entropy) in each MAF bin, and the shadowed areas represent the quantiles [0.0, 0.01, 0.25, 0.75, 0.99, 1.0]. For the sake of readability, the envelopes for the quantiles are shown only for the first and the last cycles. We observe the strongest improvement in genotyping accuracy for variants with *MAF ≥* 0.3 and through the first cycles (2 and 3). The increase in accuracy is more limited in later cycles, as the model converges to imputation results consistent with the pool observations. The irregularities of the median line and of the envelopes, for instance around *MAF* ≃ 0.22, are due to the sparsity of markers in some parts of the MAF spectrum. Subfigure 2a: Concordance computed for all genotypes. Subfigure 2b: Cross-entropy computed for all genotypes. Note that only values between 0.0 and 1.5 cross-entropy are shown.

In Fig. 3, 4, and 5, we present the changes in the genotyping accuracy in early (N=1 to N=2), intermediate (N=11 to N=12), and late (N=41 to N=42) consecutive cycles. The improvement is computed as the difference between the accuracy in the cycles N+1 and N. In these figures, we distinguish the gain in accuracy between consecutive cycles for the set of genotype entries whose priors were updated by *repool* on the one hand, and the set of genotype priors that have remained unchanged on the other hand. What set of decoded entries are corrected might vary at each cycle. The priors passed as input to *prophaser* are considered as unchanged if the element-wise difference between the genotype probabilities in cycle N and cycle N+1 is less than *ϵ* = 10^*−*5^. We observe that the markers with updated priors have the largest increase in genotyping accuracy in all cycles and this increase is curbed through the iterations. For instance, the average concordance of markers with changed prior genotype probabilities rises by 0.03157 between the first and the second cycle, and only 0.00064 between the second last and the last cycle. This confirms that the corrections we apply have an actual and positive impact, which is moreover the strongest in the earliest cycles. In Fig. 3, we also note a gain in accuracy for the set of unchanged genotypes that are in the MAF range [0.3, 0.5]. This suggests that corrections applied to some genotypes can improve the predictions at unchanged entries, typically in the case of high LD between markers.

**Fig 3.**
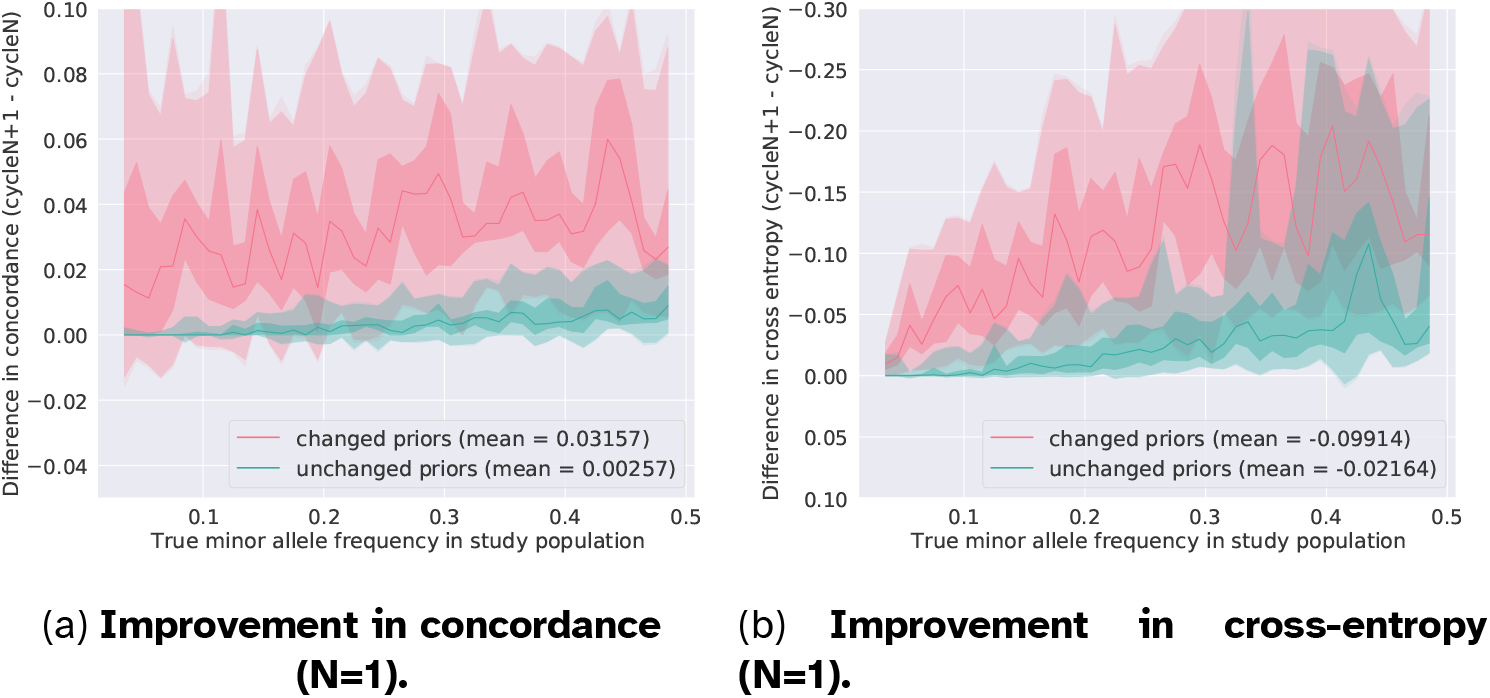
Improvement in genotyping accuracy between cycle 2 and cycle 1 (N=1) computed separately for the genotypes with an *updated* prior and for the genotypes with an *unchanged* prior (dampening factor *w* = 0.01). The plain line shows the median accuracy score in each bin. The shadowed areas display the quantiles [0.0, 0.01, 0.25, 0.75, 0.99, 1.0]. The largest gain in accuracy is obtained for the set of genotypes whose priors were corrected, but an additional round of imputation also improves, to a smaller extent, the genotyping accuracy for the predicted genotypes that were consistent with the pooled outcomes and that were therefore not changed. In both sets of priors, the greatest improvements are observed for the common variants. The genotyping accuracy of rare variants is slightly improved as well in the case where the priors are updated. Subfigure 3a: Positive values indicate that the concordance achieved after the cycle N+1 is higher than the concordance after cycle N, that is, the iteration N+1 has improved the genotyping accuracy. Subfigure 3b: Negative values indicate that the cross-entropy achieved after the cycle N+1 is lower than the concordance after cycle N, that is, the iteration N+1 has improved the genotyping accuracy. The y-axis is reverted in order to display the improvements above the origin.

**Fig 4.**
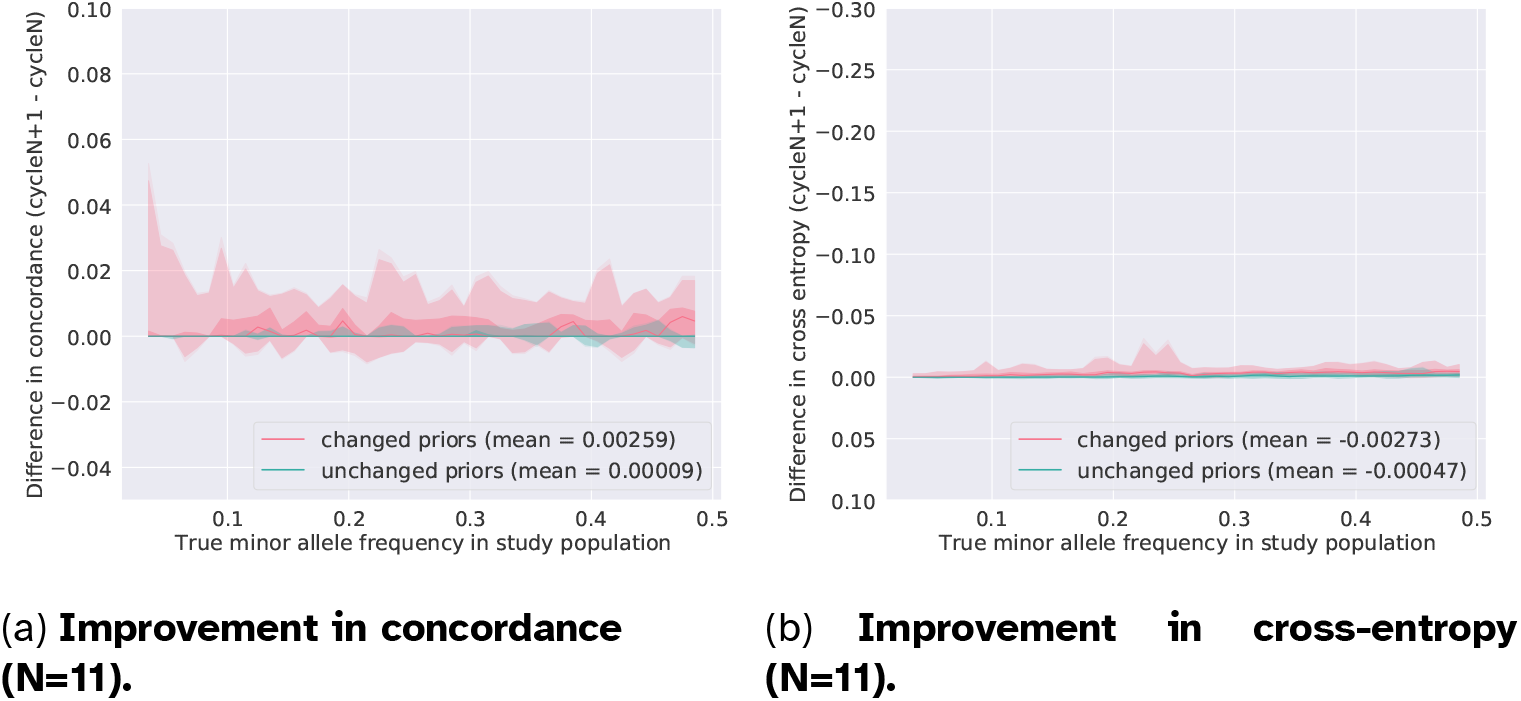
Improvement in genotyping accuracy between cycle 12 and cycle 11 (N=11) computed separately for the genotypes with an *updated* prior and for the genotypes with an *unchanged* prior (dampening factor *w* = 0.01). The plain line shows the median accuracy score in each bin. The shadowed areas display the quantiles [0.0, 0.01, 0.25, 0.75, 0.99, 1.0]. We still observe small gain in accuracy for the genotypes which have corrected priors, on the contrary to the markers with unchanged priors that seem to have reached optimal values for the predicted genotypes. Subfigure 4a: The pink envelope in a positive range suggests that changing some priors still triggers swaps to correct predictions. Subfigure 4b: Cross-entropy is barely modified, which indicates that in spite of changes in favor of the correct true genotype, the predictions are weak i.e. the largest genotype probability is close to 0.5.

**Fig 5.**
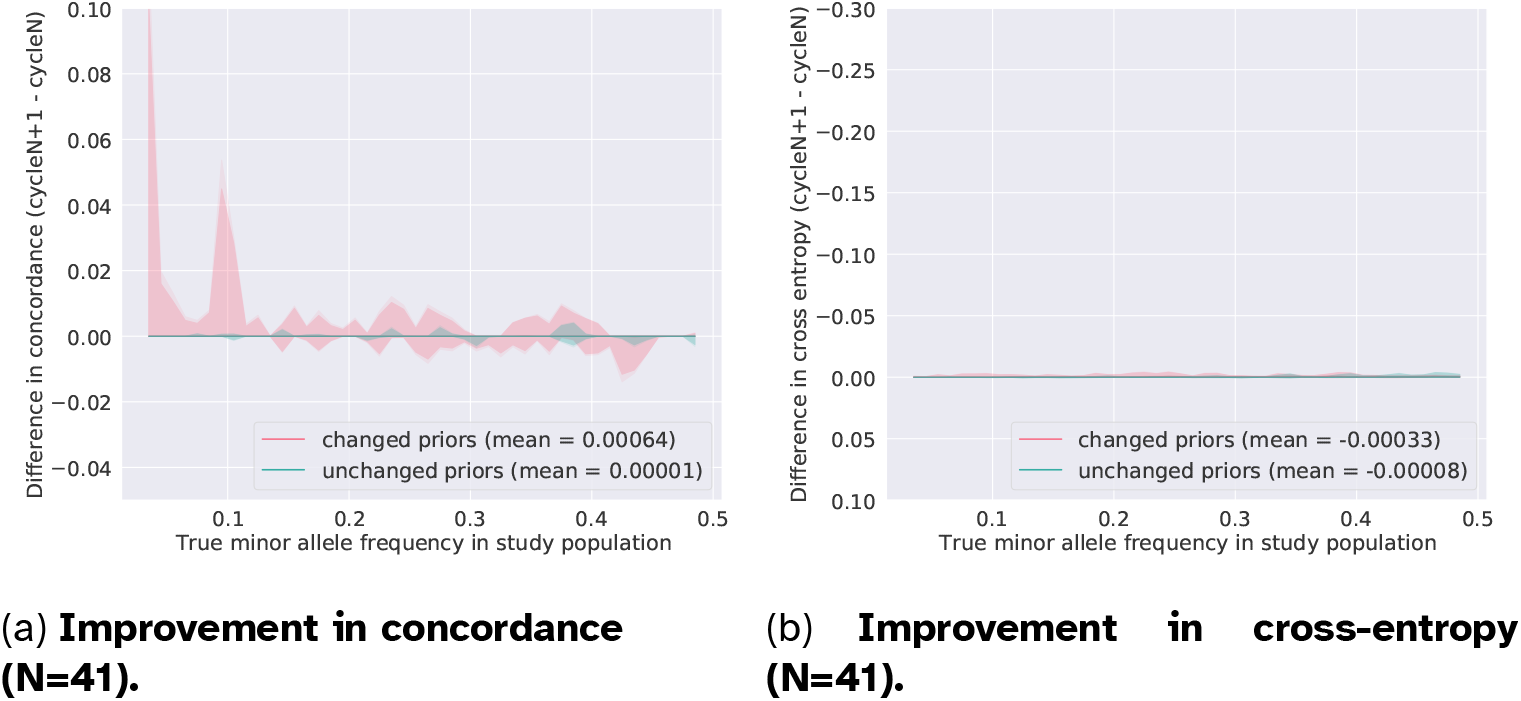
Improvement in genotyping accuracy between cycle 42 and cycle 41 (N=41) computed separately for the genotypes with an *updated* prior and for the genotypes with an *unchanged* prior (dampening factor *w* = .01). The plain line shows the median accuracy score in each bin. The shadowed areas display the quantiles [0.0, 0.01, 0.25, 0.75, 0.99, 1.0]. We no longer observe any significant gain in accuracy that could be correlated to changes in the prior genotype probabilities (on average, concordance is improved by 0.00064). Subfigure 5a: We note large positive peaks around *MAF* = 0.1 and *MAF* = 0.02 and a smaller negative peak around *MAF* = 0.42. These peaks suggest a sort of instability in the results, likely due to weak genotype predictions. In other words, at some loci and for some individuals, both homozygous genotypes might have close imputed probability. Subfigure 5b: The cross-entropy score is slightly modified, which also points in favor of almost equally likely probabilities for the opposite homozygotes. The tables below show an example of a true genotype value that is changed to the incorrect homozygote in cycle 42 (detrimental swap), and an example of a true genotype value that is changed correctly (favorable swap). We do observe homozygous genotype probabilities that are close to 0.5.

Table 2 tracks the changes in the imputed probability in the pool with the samples A1060_A1060, A1075_A1075, A1394_A1394, and A1082_A1082, for the variant 1:11814388. After the first cycle of decoding and imputation, all samples, but A1060_A1060, are correctly imputed. A1060_A1060 is first predicted to the opposite homozygote and swaps to the correct prediction after 23 cycles of correction and imputation. In later cycles, this correct prediction gets stronger since the posterior probability of the homozygous genotype rises from 50.9% after the cycle 23 to 69.6% after the cycle 42. That is, the cross-entropy for this genotype drops through the cycles. The example of A1060_A1060 demonstrates that our method can continuously and consistently improve the posterior genotype probabilities. A closer analysis of the decoded genotype likelihoods in Table S1.1. reveals that the successive corrections increase the likelihood for the allele 1 in the sample A1060_A1060. For the sample A1082_A1082, we observe that the posterior genotype probabilities oscillate before converging to the correct prediction. In order to understand this behavior, we need to consider the data in the overlapping pool in the different cycles. In short, the overlapping pool increases the likelihood for allele 0, due to an inconsistency in its absence in that pool, resulting in a competing effect on A1082_A1082. The weakening in the correct genotype prediction for the samples A1075_A1075 and A1394_A1394 are explained by the successive adjustments that raise the likelihood of genotypes carrying the alternate allele. Nonetheless, the imputed genotypes remain correct.

**Table 2.** Examples of genotypes predicted for 4 samples in a pool at marker 1:11814388 through several cycles of pooling and imputation (*w* = 0.01).

|  | A1060_A1060 |  |  | A1075_A1075 |  |  | A1394_A1394 |  |  | A1082_A1082 |  |  |
| --- | --- | --- | --- | --- | --- | --- | --- | --- | --- | --- | --- | --- |
| True genotypes | 0.0 | 0.0 | 1.0 | 1.0 | 0.0 | 0.0 | 1.0 | 0.0 | 0.0 | 1.0 | 0.0 | 0.0 |
| Cycle 1 |  |  |  |  |  |  |  |  |  |  |  |  |
| Imputed genotypes | 0.99989 | 0.0 | 0.00011 | 1.0 | 0.0 | 0.0 | 0.999892 | 0.0 | 0.000108 | 0.489336 | 0.0 | 0.510664 |
| Cycle 2 |  |  |  |  |  |  |  |  |  |  |  |  |
| Imputed genotypes | 0.999824 | 0.0 | 0.000176 | 1.0 | 0.0 | 0.0 | 0.999828 | 0.0 | 0.000172 | 0.657397 | 0.0 | 0.342603 |
| Cycle 11 |  |  |  |  |  |  |  |  |  |  |  |  |
| Imputed genotypes | 0.938863 | 0.0 | 0.061137 | 0.999998 | 0.0 | 0.000002 | 0.954346 | 0.0 | 0.045654 | 0.50544 | 0.0 | 0.49456 |
| Cycle 12 |  |  |  |  |  |  |  |  |  |  |  |  |
| Imputed genotypes | 0.907764 | 0.0 | 0.092236 | 0.999998 | 0.0 | 0.000002 | 0.933557 | 0.0 | 0.066443 | 0.493936 | 0.0 | 0.506064 |
| Cycle 22 |  |  |  |  |  |  |  |  |  |  |  |  |
| Imputed genotypes | 0.514556 | 0.0 | 0.485444 | 0.999971 | 0.0 | 0.000029 | 0.734207 | 0.0 | 0.265793 | 0.575863 | 0.0 | 0.424137 |
| Cycle 23 |  |  |  |  |  |  |  |  |  |  |  |  |
| Imputed genotypes | 0.491217 | 0.0 | 0.508783 | 0.999966 | 0.0 | 0.000034 | 0.728201 | 0.0 | 0.271799 | 0.587209 | 0.0 | 0.412791 |
| Cycle 41 |  |  |  |  |  |  |  |  |  |  |  |  |
| Imputed genotypes | 0.307134 | 0.0 | 0.692866 | 0.999513 | 0.0 | 0.000487 | 0.705797 | 0.0 | 0.294203 | 0.69339 | 0.0 | 0.30661 |
| Cycle 42 |  |  |  |  |  |  |  |  |  |  |  |  |
| Imputed genotypes | 0.304208 | 0.0 | 0.695792 | 0.999441 | 0.0 | 0.000559 | 0.704606 | 0.0 | 0.295394 | 0.696172 | 0.0 | 0.303828 |

True genotypes and imputed genotypes are given as genotypes probabilities. Red and green highlighting shades indicates how much wrong or correct the predicted genotype is with respect to the true one.
For the sample A1060\_A1060, rectification to the correct genotype occurs after the cycle 23 has completed.

In Fig. 5, we also observe sharp peaks in the concordance of set of markers with changed priors, whereas the cross-entropy is uniform and almost equal to 0. We interpret those peaks in the last cycle as “favorable” and “detrimental” genotype swaps that describe the cases where an incorrect prediction after the cycle 41 becomes correct after the cycle 42, and conversely. Such swaps also suggest some instability of a few predictions that are either weakly correct or weakly wrong, that is the probabilities for both homozygous genotypes are close to 0.5. Weak predictions in favor of the opposite homozygotes at a given entry have a more noticeable impact on the concordance than on the cross-entropy since the difference in concordance is 1 whereas the order of magnitude of the difference in cross-entropy is 10^*−*1^. For instance, a change in the genotype probability from *Pr*(*G* = 2) = 0.479 to *Pr*(*G* = 2) = 0.505 in the example of favorable swap results in an obvious peak down, but has a little impact on the cross-entropy. On the contrary, a change with a similar amplitude in a denser MAF interval, such as the example of detrimental swap, is less noticeable.

Table 3 gives more detailed counts of the genotypes that are corrected between consecutive cycles. The counts across all samples are aggregated within MAF intervals, such that we find that for the rare variants, about 2100 genotypes are updated in each cycle, that is ca. 5.6% of the 37696 data points in the interval. This low percentage suggests that the posterior genotypes after imputation are mostly consistent with the pooled observations. The MAF intervals that span over 0.2 to 0.5 have the largest shares of corrected genotypes with a range from ca. 32% up to nearly 40%. The counts of corrected genotypes slightly vary through the cycles, for instance, in the MAF interval [0.4, 0.5], 35.9% of the genotypes are updated between the cycles 1 and 2, and 39.8% between the cycles 41 and 42. In contrast, the MAF interval [0.05, 0.1] shows the opposite trend with 13.4% of corrected genotypes from the cycle 1 to 2, and eventually 11.9% from the cycle 41 to 42. Across all variants and samples, about 25%of the data points are updated after each cycle.

**Table 3.**
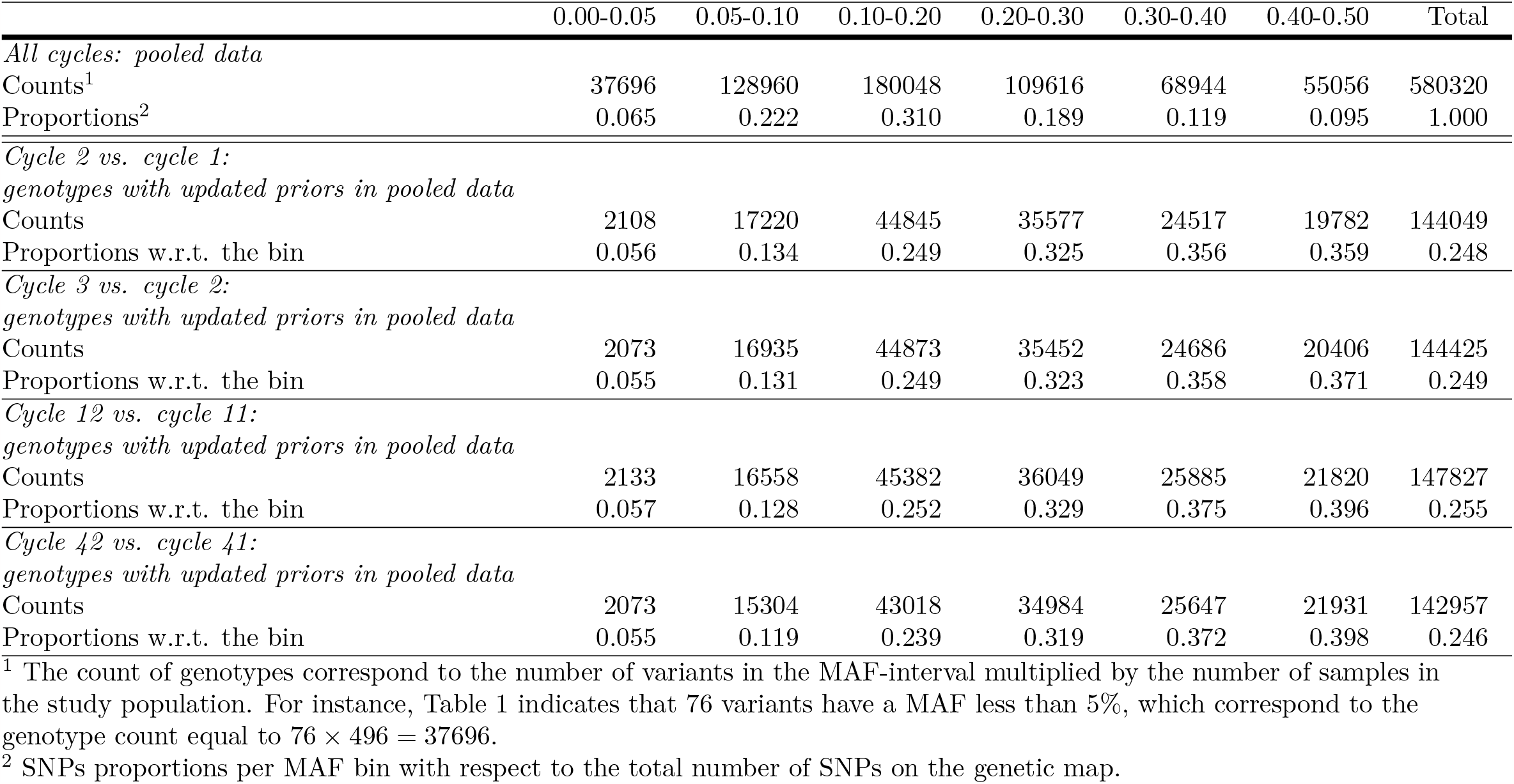
Statistics for genotypes (variants x samples) per MAF bin in the population of inbred lines (pooled data, dampening factor for the pooled genotype likelihoods equal to 0.01)

|  | 0.00-0.05 | 0.05-0.10 | 0.10-0.20 | 0.20-0.30 | 0.30-0.40 | 0.40-0.50 | Total |
| --- | --- | --- | --- | --- | --- | --- | --- |
| *All cycles: pooled data* |  |  |  |  |  |  |  |
| Counts<sup>1</sup> | 37696 | 128960 | 180048 | 109616 | 68944 | 55056 | 580320 |
| Proportions<sup>2</sup> | 0.065 | 0.222 | 0.310 | 0.189 | 0.119 | 0.095 | 1.000 |
| *Cycle 2 vs. cycle 1:*<br>*genotypes with updated priors in pooled data* |  |  |  |  |  |  |  |
| Counts | 2108 | 17220 | 44845 | 35577 | 24517 | 19782 | 144049 |
| Proportions w.r.t. the bin | 0.056 | 0.134 | 0.249 | 0.325 | 0.356 | 0.359 | 0.248 |
| *Cycle 3 vs. cycle 2:*<br>*genotypes with updated priors in pooled data* |  |  |  |  |  |  |  |
| Counts | 2073 | 16935 | 44873 | 35452 | 24686 | 20406 | 144425 |
| Proportions w.r.t. the bin | 0.055 | 0.131 | 0.249 | 0.323 | 0.358 | 0.371 | 0.249 |
| *Cycle 12 vs. cycle 11:*<br>*genotypes with updated priors in pooled data* |  |  |  |  |  |  |  |
| Counts | 2133 | 16558 | 45382 | 36049 | 25885 | 21820 | 147827 |
| Proportions w.r.t. the bin | 0.057 | 0.128 | 0.252 | 0.329 | 0.375 | 0.396 | 0.255 |
| *Cycle 42 vs. cycle 41:*<br>*genotypes with updated priors in pooled data* |  |  |  |  |  |  |  |
| Counts | 2073 | 15304 | 43018 | 34984 | 25647 | 21931 | 142957 |
| Proportions w.r.t. the bin | 0.055 | 0.119 | 0.239 | 0.319 | 0.372 | 0.398 | 0.246 |

<sup>1</sup> The count of genotypes correspond to the number of variants in the MAF-interval multiplied by the number of samples in the study population. For instance, Table 1 indicates that 76 variants have a MAF less than 5%, which correspond to the genotype count equal to $76 \times 496 = 37696$ . <sup>2</sup> SNPs proportions per MAF bin with respect to the total number of SNPs on the genetic map.

The overall stability in the number of corrected genotypes after each marker observed in Table 3 might look discordant with the results in Fig. 2, where the gain in accuracy after correction and imputation decreases through the cycles. Therefore, we calculate the total correction that is added to the pooled decoded data in each cycle, and we observe that the total correction in the decoded genotypes drops through the cycles (supplementary Fig. S2.10). That is, although the number of genotypes that are corrected is approximately the same in any cycle, the amplitude of the correction per genotype decreases strongly over the iterations.

These gradually smaller corrections, as well as the trend in Fig. 2 and 3-5, indicate a convergence of our coupled model.

### Computational performance of the coupled iterations

In any cycle *N >* 1, correcting the decoded data with *repool* takes about 7 seconds, with a very modest memory usage. The computational performance of imputation with *prophaser* is similar to the cycle 1, that is about 475 milliseconds per study sample, and require an insignificant amount of memory. Hence, from the perspective of the computational complexity, the gains in genotyping accuracy achieved thanks to further cycles of repooling and imputation have a very low cost.

## Discussion

We want to stress that the prior adjustment step does not claim to actually produce accurate priors for the individual genotypes. Rather, the aim is to produce a set of inputs that, when combined with the imputation methodology, produce accurate posterior probabilities. The improvements in concordance as well as cross-entropy confirm this.

We obtain accuracy levels that are in the same range than in the studies mentioned earlier [4, 5] without parameterizing the distribution of the genotypes based on the Hardy-Weinberg equilibrium (HWE) as in [4]. Both *repool* and *prophaser* could handle various levels of heterozygosity in the population. They implicitly model a general diploid population, with a high frequency of heterozygotes, but based on the properties of the observed data, they respect the inbred nature of the processed input population in our dataset. Such flexibility mainly lies in the likelihood framework in which the genotype data are expressed. Moreover, the pedigree-free implementation allows for using our strategy in various types of populations and breeding schemes.

The gain in genotyping accuracy achieved through additional cycles indicates that the feedback structure successfully combines the strengths of imputation and pooling, which eventually yields very precise predictions for the genotypes at any variant. The improvements at each cycle confirm that the decoding and the imputation models exploit complementary sources of information, that are the combinatorial constraints local to the pools on the one hand, on the other hand the inter- and intra-individual genetic structure. We interpret the convergence of our coupling strategy as each information source is progressively exhausted through the iterations. As in other iterative methods, we believe that exploring adaptive values of the dampening factor *w* could contribute to optimize the trade-off between the gain in accuracy and the number of cycles to execute.

Regardless of the number of cycles executed, our strategy has the advantage of not requiring any extra sequencing nor testing on microarrays. Instead, we exclusively rely on array data and on the global genetic information carried in the library of reference haplotypes. The computational cost of each additional iteration is mainly driven by the size of the reference panel in the imputation step. That is, with a small reference panel such as the one we have, the time complexity remains very low and executing the imputation on CPU resources only demonstrates sufficient performance. In the case larger reference panels should be investigated, we have also implemented a version of *prophaser* that is suitable for execution on GPU.

## Abbreviations

GS: genomic selection;
GWAS: genome wide association studies;
LD: linkage disequilibrium;
MAF: minor allele frequency;
MAGIC: multiparent advanced generation intercross;
NGS: next generation sequencing;
NIAB: National Institute of Applied Botany;
SNP: single nucleotide polymorphism.

## Supporting information

**S1 File. repool: A decoding algorithm for iterative adjustment of prior probabilities from pool experiments based on feedback from imputation**. Detailed example of calculations for applying point-wise correction of the decoded genotype likelihoods.

**S2 File. Supplemental figures and tables**. Supplementary figures and tables for additional results with other dampening factors.

## Acknowledgments

The computing resources were provided by National Academic Infrastructure for Supercomputing in Sweden (NAISS, formerly SNIC) through Uppsala Multidisciplinary Center for Advanced Computational Science (UPPMAX) under Project SNIC 2022/22-697.

## Optional sections

### Authors’ contributions

CN conceived the study and implemented the code for point-wise correction of the decoded genotype likelihoods. CC developed the script for executing the cycles as well as the code for generating the graphics, ran the experiments, and wrote the first draft of the manuscript. All authors edited the manuscript and contributed to analyzing the results as well as to the conclusions. All authors read and approved the final manuscript.

### Funding

Research project funded by Formas, The Swedish government research council for sustainable development. Grant Nr. 2017-00453. Cost-effective genotyping in plant and animal breeding using computational analysis of pooled samples.

### Ethics approval

Not applicable.

### Consent to participate

Not applicable.

### Consent for publication

Not applicable.

### Availability of data and materials

The datasets supporting the conclusions of this article are available at http://mtweb.cs.ucl.ac.uk/mus/www/MAGICdiverse/MAGICdiverseFILES/ (genotype data) and https://urgi.versailles.inra.fr/download/iwgsc/IWGSCRefSeqAnnotations/v1.0/iwgscrefseqv1.0recombinationrateanalysis.zip (genetic maps).

We refer to a a workflow which can be found at https://github.com/camcl/poolimputeSNPs/tree/iterpoolimp for reproducing our results and simulations. Note however that the cycles of correcting decoding and imputation were run with the computational resources provided by UPPMAX which uses Slurm and specific settings. This implies some (extensive) refactoring of our shell scripts. This workflow uses source codes that are accessible at https://github.com/camcl/genotypooler/tree/iterpoolimp (*genotypooler*) and at https://github.com/scicompuu/prophaser/tree/multilevel (*prophaser*).

---

Example of detrimental last swap in genotype prediction with nearly equally likely GP for the opposite homozygotes:

Sample A1343_A1343 at variant 1:471263300 (*MAF* = 0.429435) has GT = 0/0, which is correctly predicted after cycle 41 but is predicted to 1/1 in cycle 42.

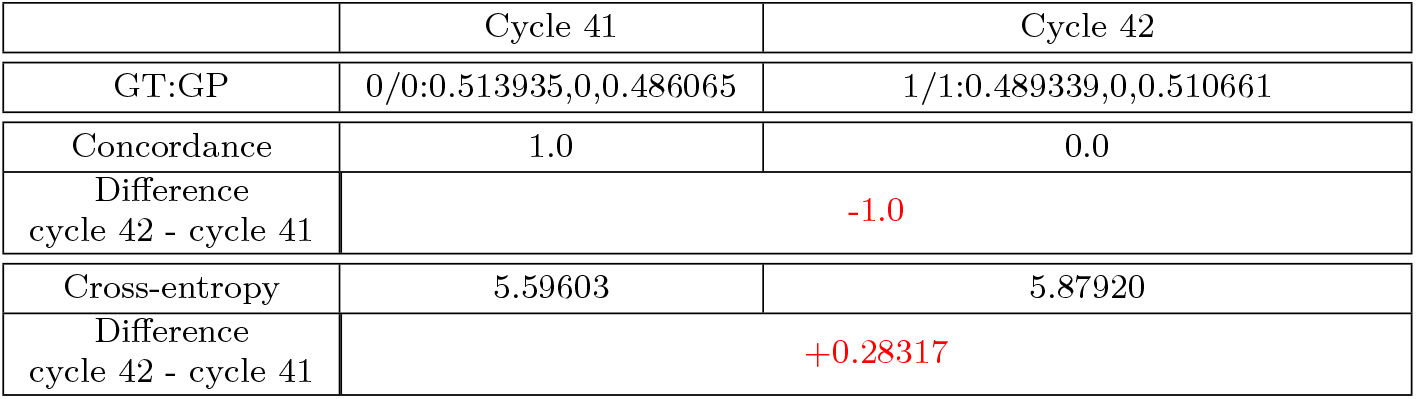

Example of favorable last swap in genotype prediction with nearly equally likely GP for the opposite homozygotes:

Sample A1118_A1118 at variant 1:29607405 (*MAF* = 0.032258) has GT = 1/1, which is incorrectly predicted to 0/0 after cycle 41 but rectified to 1/1 in cycle 42.

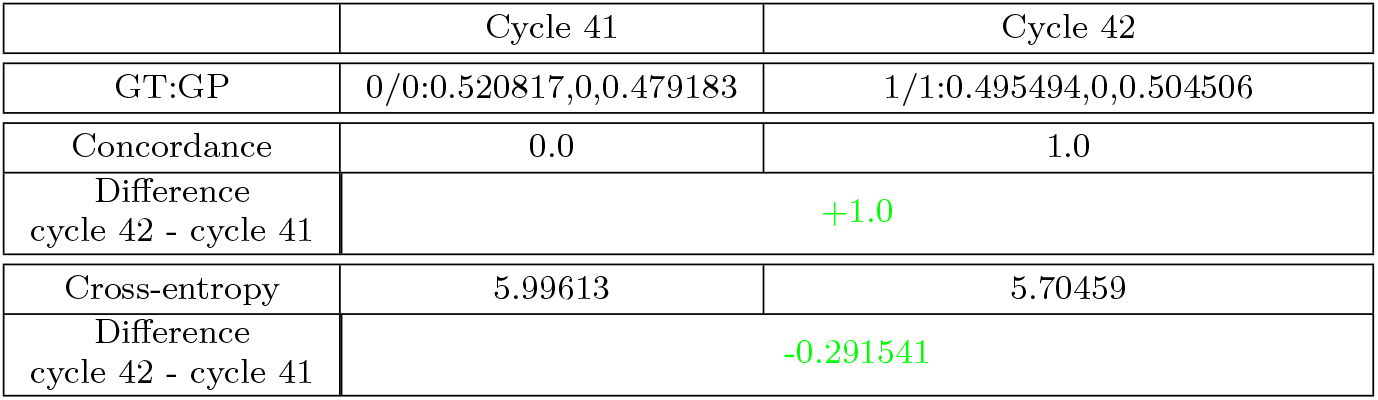

